# Discovery of CMX990: A Potent SARS-CoV-2 3CL Protease Inhibitor Bearing a Novel Covalent Warhead

**DOI:** 10.1101/2023.10.24.563688

**Authors:** N. G. R. Dayan Elshan, Karen C. Wolff, Laura Riva, Ashley K. Woods, Gennadii Grabovyi, Katy Wilson, Alireza Rahimi, James Pedroarena, Sourav Ghorai, Anil Kumar Gupta, Armen Nazarian, Frank Weiss, Yuyin Liu, Wrickban Mazumdar, Lirui Song, Neechi Okwor, Jacqueline Malvin, Malina A. Bakowski, Nathan Beutler, Melanie G. Kirkpatrick, Amal Gebara-Lamb, Edward Huang, Van Nguyen-Tran, Victor Chi, Shuangwei Li, Thomas F. Rogers, Case W. McNamara, Jian Jeffrey Chen, Sean B. Joseph, Peter G. Schultz, Arnab K. Chatterjee

## Abstract

There remains a need to develop novel SARS-CoV-2 therapeutic options that improve upon existing therapies by increased robustness of response, fewer safety liabilities, and global-ready accessibility. Functionally critical viral main protease (M^pro^, 3CL^pro^) of SARS-CoV-2 is an attractive target due to its homology within the coronaviral family, and lack thereof towards human proteases. In this disclosure, we outline the advent of a novel SARS-CoV-2 3CL^pro^ inhibitor, CMX990, bearing an unprecedented trifluoromethoxymethyl ketone warhead. Compared with the marketed drug nirmatrelvir (combination with ritonavir = Paxlovid^TM^), CMX990 has distinctly differentiated potency (∼5x more potent in primary cells) and human *in vitro* clearance (>4x better microsomal clearance and >10x better hepatocyte clearance), with good *in vitro*-*in vivo* correlation. Based on its compelling preclinical profile and projected once or twice a day dosing supporting unboosted oral therapy in humans, CMX990 advanced to a Phase 1 clinical trial as an oral drug candidate for SARS-CoV-2.

## INTRODUCTION

With ∼770 million confirmed cases and ∼7 million deaths worldwide to date, the SARS-CoV-2 pandemic has had a resounding impact around the globe.^1^ While the advent of prophylactic vaccines has helped overcome pandemic status,^2^ many health hurdles still remain for those who contract the disease and/or are not vaccinated due to personal, socioeconomic, geographic, or other adverse circumstances. In account of this, and to prepare for future pandemics, many drug discovery programs were pursued around the world for post-infection treatment options. Some of the more successful outcomes from such efforts include the drugs remdesivir (Veklury®, Gilead Sciences), nirmatrelvir (+ ritonavir = Paxlovid®, Pfizer Inc.), and ensitrelvir (Xocova®, Shionogi & Co.).^3–5^ Although having shown some level of efficacy in clinical trials, each of these therapies has limitations in areas such as robustness of response, therapeutic index and toxicology, drug-drug interaction (DDI) concerns, affordability, and wide-spread availability. Therefore, there remains a major interest and need to develop alternative therapies for coronaviral infections. Herein, we describe a program that led to the development of a unique SARS-CoV-2 drug candidate that reached Phase 1 clinical trials.

At the onset of the coronavirus pandemic in 2020, Calibr leveraged its drug repurposing library ReFRAME (Repurposing, Focused Rescue, and Accelerated Medchem),^6^ both internally and with external collaborators, to find potential hits that could be quickly adapted as effective antivirals to combat SARS-CoV-2. Via these screens, multiple clinical-stage cysteine/serine protease inhibitors such as emricasan, VBY-825, and dutacatib (**Figure 1A**), were found to have moderate SARS-CoV-2 antiviral potency. This made sense as viral 3CL^pro^ is a main enzyme involved in SARS-CoV-2 replication, and there is significant homology across various proteases.^7^ SARS-CoV-2, belonging to the genus Betacoronavirus, encodes four structural proteins (spike, envelope, membrane, and nucleocapsid) and several accessory proteins. Two polyproteins (pp1a/pp1ab) must be cleaved into their individual, nonstructural proteins for successful viral replication.^8^ Two viral proteases are essential and responsible for processing the polyproteins: the 3C-like protease (3CL protease, 3CL^pro^, CL^pro^; also referred to as “main protease”, M^pro^) and a papain-like protease (PL^pro^).^9^

**Figure 1:**
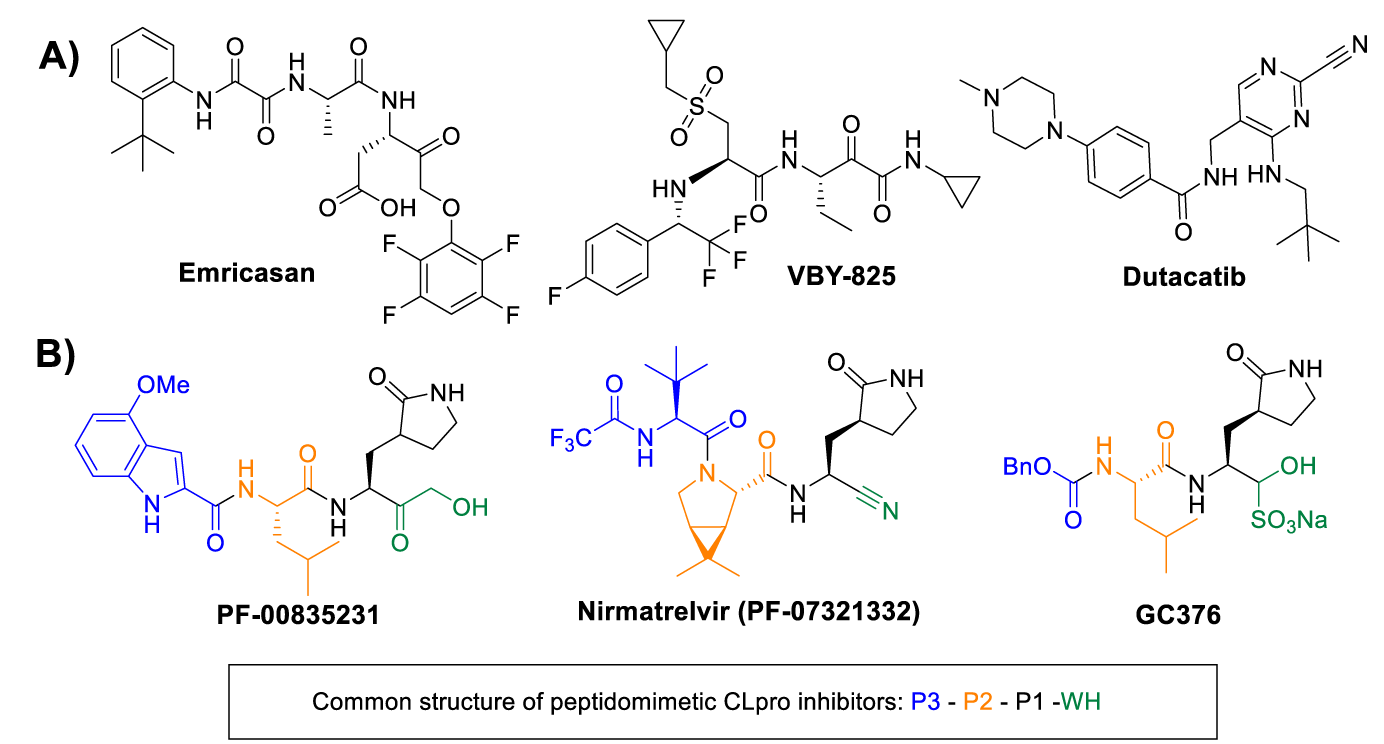
Representative known protease inhibitors. A) Initial ReFRAME library hits identified as moderate (low µM) potency inhibitors of SARS-CoV-2. B) Peptidomimetic SARS-CoV-2 CL^pro^ inhibitors PF-00835231 (Pfizer), PF-07321332 (Pfizer), and GC376 (Anivive). Generally, up to four distinct structural regions are found in all peptidomimetic CL^pro^ inhibitors: warhead (WH), P1 (amino acid 1, recognition element with a lactam side chain), P2 (AA 2), and P3 (a *N*-Cap, or a third AA). Common warheads include hydroxymethyl ketones (e.g., PF-00835231), nitriles (e.g., nirmatrelvir), aldehydes or bisulfite adducts thereof (e.g., GC376), and aryloxymethyl ketones (e.g., emricasan).

CL^pro^ cleaves polypeptides after a glutamine residue in the P1 position of the substrate, which is a unique activity not observed in other human proteases and suggests that this viral protease can be specifically and selectively inhibited by a small-molecule inhibitor.^10^ The homology between coronaviral main proteases provides an attractive avenue to develop therapeutics that could also be capable of tackling future threats from mutated viral variants as well as other genera of coronaviruses. CL^pro^ has four active sites (S1′, S1, S2, and S4), which can be accommodated by distinct structural regions of a peptidomimetic substrate/inhibitor amenable for structure-activity relationship (SAR) optimization (**Figure 1B**): A warhead (WH), P1 (amino acid [AA] 1, recognition element with a lactam side chain), P2 (AA 2), and P3 (a *N*-Cap, or a third AA). The warhead that engages the catalytic site Cys145 plays an important role in the potency of such inhibitors, given that Cys145 is an essential residue for antiviral activity.^11^ In addition, it is well established that a glutamine derivative (γ-lactam) is highly preferred to occupy the S1 site of cysteine proteases, which not only mimics the P1 glutamine of the native substrates but also increases the activity of inhibitors.^12^ Many of these characteristics about the engagement of coronaviral CL^pro^ were well established in the seminal work done by Pfizer Inc. and others during the SARS-CoV-1 outbreak, leading to CL^pro^ inhibitor drug candidates such as PF-00835231,^13^ which was the active form of one of the very first candidates going into Phase 1 clinical trials for SARS-CoV-2 treatment.

## RESULTS AND DISCUSSION

Extending our ReFRAME starting points via incorporation of literature knowledge of protease inhibitor designs (including caspase/cathepsin literature), we have explored SAR across all regions of this peptidomimetic target template, leading to a library of over 1600 compounds to date.^14^ Only a key SAR trajectory (**Figure 2**) and the performance characteristics of selected representative compounds (**Table 1**) leading to the clinical candidate **CMX990** are highlighted in this disclosure. The SAR evaluations happened across both P2-non-prolines as well as P2-proline derivatives. Key optimization parameters during early development were to improve the potency and human microsomal clearance of PF-00835231. The warhead change from the hydroxymethyl ketone in PF-00835231 to an aryloxymethyl ketone (**1**) led to a ∼9-fold increase in potency (**Figure 2**). However, the microsomal clearance of this warhead was rapid, leading to poor PK profiles. Opening of the indole ring to get the aryl oxamide analog **2** led to a modest increase in potency, but this made the microsomal clearance worse. A number of additional warheads were screened including but not limited to, nitriles, ketoamides, aldehydes, and prodrugs of hydroxymethyl ketone. All these warheads either lacked the potency and/or the clearance (*in vitro* / *in vivo*) profile we were looking for. We therefore decided to develop a novel covalent warhead for this compound series, resulting in the trifluoromethoxymethyl ketone warhead compound **3**. Compared to the preceding aryloxymethyl ketone analog **2**, the new warhead analog **3**, resulted in >3x improved potency (HeLa-ACE2 CoV2 EC_90_ = 17 nM) and >30x improved *in vitro* clearance. Encouraged by this breakthrough finding, we incorporated the novel warhead across a wide array of P3-P2 combinations, where incorporation of a proline at P2 gave the early lead **4**, which retained much of the potency gained in the non-proline analog **3**. This lead also had far superior oral bioavailability in rodent and dog pharmacokinetic studies (**Table 1**), which was expected given the known ability of proline incorporation to enhance the exposure of orally administered peptidomimetics.^15^ Aryl oxamide containing compounds (f.ex. **3**) were unstable in rat/mouse plasma, so we used hamster as the rodent species for pharmacokinetic studies of such analogs (**Table 1**). Despite the early interest, we discontinued development of **4** and other aniline oxamide leads, due to a drug-related mutagenicity response seen in an Ames assessment.

**Figure 2:**
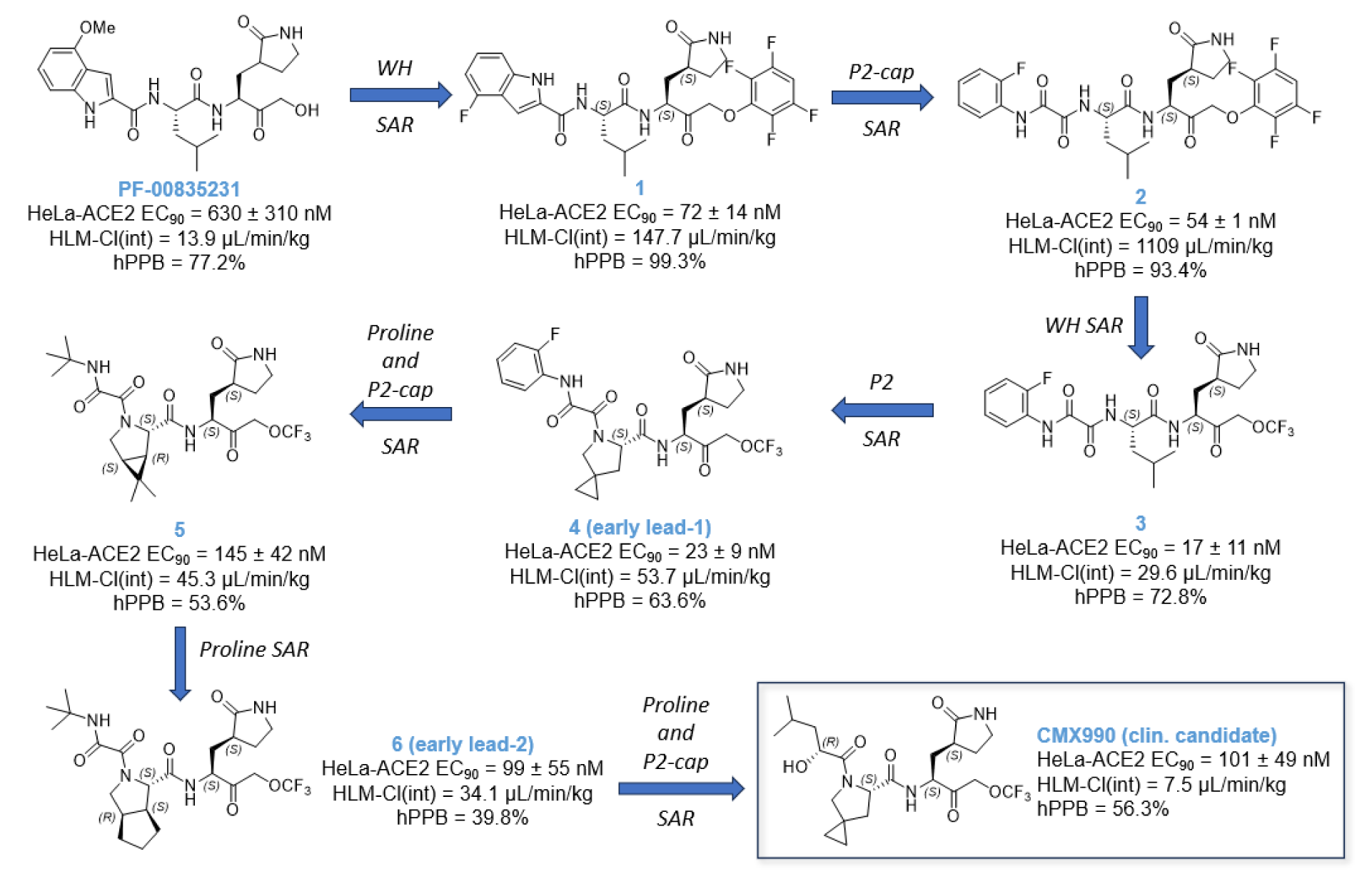
Representative SAR summary trajectory to early program leads, including clinical candidate CMX990. SAR across all regions of the compounds were carried out in parallel, leading to a series of >1600 total compounds to date. WH = warhead.

**Table 1:** Representative compounds and profiling data for CMX990 series SAR.

| Compound | logD | CoV-2<br>IC<sub>50</sub><br>(nM) | CoV-2 EC<sub>90</sub><br>(nM) |  | hPPB (%) | HLM<br>Cl<sub>int</sub><br>(μL/min/mg) | IV PK Clp<br>(mL/min/kg) |  | F(PO) % |  |
| --- | --- | --- | --- | --- | --- | --- | --- | --- | --- | --- |
|  |  |  | ALI-HBEC | HeLa-ACE2 |  |  | Mu | Dog | Mu | Dog |
| 1 | 4.4 | n.d. | n.d. | 72 | 99.3 | 147.7 | 31.8<sup>a</sup> | - | < 1<sup>a</sup> | - |
| 2 | 3.3 | 14.8 | n.d. | 54 | 93.4 | 1109.4 | 47.6<sup>a,b</sup> | 50.3 | 1.3<sup>a,b</sup> | 1.5 |
| 3 | 2.8 | 76.3 | 133 | 17 | 72.8 | 29.6 | 173<sup>a</sup> | 53.4 | < 1<sup>a</sup> | 21.5 |
| 4 | 2.4 | 16.1 | 14.3 | 23 | 63.6 | 53.7 | 129 | 28.7 | 6.3 | 38.0 |
| 5 | 1.6 | 81.1 | 125 | 145 | 53.6 | 45.3 | 65.9 | 36.9 | 32.4 | 23.4 |
| 6 | 2.0 | 28.8 | 19.3 | 99 | 39.8 | 34.2 | 45.1 | 38.8 | 25.2 | 21.6 |
| 7 | 2.6 | 24.0 | n.d. | 117 | 54.8 | 62.2 | - | 36.2 | - | - |
| 8 | 3.0 | 34.7 | n.d. | 61 | 43.5 | 26.2 | 53.2<sup>c</sup> | - | 51.3<sup>c</sup> | - |
| 9 | 2.0 | 36.5 | n.d. | 108 | 58.8 | 16.3 | - | 40.3 | - | - |
| 10 | 1.3 | 35.4 | n.d. | 346 | < 1 | 12.8 | 71.0 | 11.7 | 7.2 | 28.0 |
| 11 | 2.0 | 34.5 | 50.8 | 62 | 31.7 | 8.4 | 35.6<sup>c</sup> | 51.8 | 35.3<sup>c</sup> | 35.7 |
| 12 | 1.0 | 33.7 | 20.5 | 24 | 47.3 | 17.4 | 11.1<sup>c</sup> | 56.9 | 20.7<sup>c</sup> | 25.0 |
| 13 | 3.6 | 34.8 | 14 | 38 | 67.5 | 19.2 | n.c. | 15.0 | 2.4 | 35.3 |
| 14 | 2.0 | 60.1 | 51.8 | 138 | 34.3 | 12.2 | 107 | 18.4 | 19.0 | 48.7 |
| 15 | 0.3 | 160 | n.d. | 245 | 51.8 | 13.0 | n.c.<sup>a</sup> | 23.3 | 4.6<sup>a</sup> | 37.7 |
| 16 | 2.6 | 22.0 | n.d. | 50 | 50.1 | 14.9 | 19.3<sup>c</sup> | 23.7 | 10.2<sup>c</sup> | 57.2 |
| **CMX990** | **2.3** | **23.4** | **9.6** | **101** | **56.3** | **7.5** | **125** | **23.0** | **14.5** | **52.8** |
| Nirmatrelvir | 1.5 | 28.1 | 44.8 | 148 | 54.8 | 21.8 | 35.8 | 1.92 | 53.5 | 68.7 |
| PF-00835231 | 1.2 | 101 | 81.4 | 630 | 77.2 | 13.9 | 86.4 | - | 1.0 | - |

*a* = hamster, *b* = data from close analog, *c* = rat
EC<sub>90</sub> = 90% effective concentration, Cl<sub>int</sub> = intrinsic clearance, Clp = plasma clearance, F = bioavailability, HLM = human liver microsomes, hPPB = human plasma protein binding, IV = intravenous, Mu = mouse, n.d. = not determined, n.c. = not calculated, PO = per os (oral)

Switching to alkyl oxamides immediately alleviated the Ames assay outcome (e.g., **5**), indicating a clear link to the aniline oxamides regarding the positive Ames signal. Additionally, moving away from the aryl oxamides helped improve the human microsomal stability, which along with further SAR of P2 proline yielded the early lead **6** that performed similarly to nirmatrelvir in many aspects (**Figure 2**, **Table 1**), including a mouse efficacy study (data not disclosed at this time).

The alkyl oxamide derivative **6** had a very favorable outlook across broad safety profiling, including a peptidase panel assessment for cross reactivity with human proteases. However, at this time nirmatrelvir was already available for treatment of SARS CoV-2, and we did not see enough differentiation from nirmatrelvir potency nor clearance profiles to have sufficient justification to advance **6** for further development. This led us to further screen P2 prolines and P3 substituents that culminated in the discovery of the clinical candidate **CMX990** (**Figures 2-3**, **Scheme 1**), which featured an α-hydroxy acid P2-cap and a 4-spiro-cycloprpyl proline at P2. Potential (though not observed in later met-ID studies) byproducts (e.g., **7** and **8; Table 1**) of manufacturing and/or metabolism of **CMX990** were characterized as well, and these retained good potency and target engagement. Alternative P3 caps screened included other forms of α-hydroxy acids such as **9**, as well as P3 truncated analogs such as **10** (**Table 1**). These modifications either compromised potency and/or had worse *in vivo* clearance. In addition to the non-proline P2 analogs in **Figure 2**, additional exciting analogs that were pursued at the time included some non-aniline-oxamide P3 compounds (f.ex. **11** and **12**), oxamide isostere derivatives such as **13**, and THF P2-cap derivative **14** (**Table 1**). These had good potency (EC_90_ = 23 – 62 nM) in HeLa-ACE2 SARS-CoV-2 assays, though **14** had a less convincing biochemical IC_50_ (>60 nM) for target engagement. Alternative warheads, such as hydroxylmethyl ketones did have advantages in clearance (2x improved HLM-Cl_int_ in **15** vs. **3**; **Table 1**) but lacked the desired potency profile (**15** HeLa-ACE2 CoV-2 EC_90_ = 245 nM, and CoV2 IC_50_ = 160 nM; **Table 1**). **CMX990** had an excellent balance of physicochemical and ADME (absorption, digestion, metabolism, and excretion) properties, allowing for high potency in target engagement, as well as favorable pharmacokinetic properties (**Tables 1-3**). It is a neutral molecule (logD_7.4_ = 1.17), with a molecular weight of 491.5 g/mol, and pH-independent good aqueous solubility, with solubility of ∼1.4-1.5 mg/mL in buffers ranging from pH 1.2-7.4. In the presence of artificial intestinal fluids near neutral pH, lower solubility was observed in fed state simulated intestinal fluid (FeSSIF) (0.734 mg/mL, pH 5.0), with higher values observed in fasted state simulated intestinal fluid (FaSSIF) (>2 mg/mL, pH 6.5).

**Figure 3:**
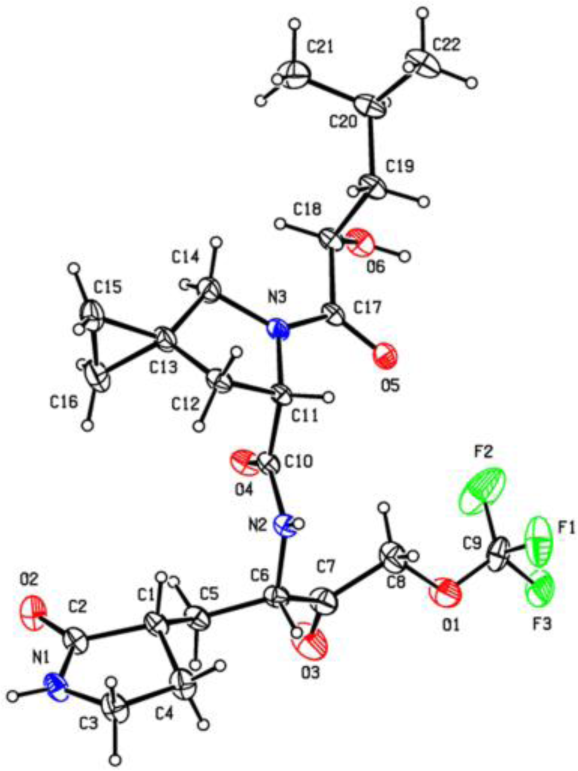
X-ray crystal structure of CMX990. Displacement ellipsoids are drawn at the 30% probability level.

The synthetic route to clinical candidate **CMX990** is illustrated in **Scheme 1**, which is also representative of the assembly of other leads with the same warhead discussed in this report. The key starting reagents are commercially available P1-ester (also used in the syntheses of analogous compounds such as nirmatrelvir) **17,** trifluoromethyl acetic acid **18**, D-leucic acid, and Boc-protected proline **21**. Alternative proline leads, such as **16** (**Table 2**) with a similar favorable potency/PK profile, were also considered for development. But the ease of access to **21** was significantly better (easily manufactured from hydroxyproline) versus these alternatives, making **CMX990** the ideal choice to progress with plans for eventual commercial-scale manufacture. The novel warhead bearing P1 amino acid was synthesized starting with the ester **17** (**Scheme 1, panel A**). Trifluoromethoxyacetic acid (**18**) is used as the key reagent to install the novel warhead on the BocNH-P1-ester **17**, yielding **19** at 47% yield. Boc deprotection of **19** with hydrochloric acid yielded the salt **20**. Using this key intermediate, **CMX990** was assembled in four linear steps starting from the Boc-proline **21** (**Scheme 1, panel B**). Following protecting group manipulation (99% yield), the proline was coupled with D-leucic acid (purchased commercially, or synthesized in house by conversion of D-Leucine using literature methods ^16^ to yield **22** (49%), which was hydrolyzed under basic conditions yielding the key carboxylic acid derivative **23**, in 93% yield. Thus obtained **23** was coupled under conventional peptide coupling conditions with the amine **20**, to give **CMX990** in as a white crystalline solid. The described process has been used to make **19** and **CMX990** in > 30 g scale. Further process optimization led to the manufacture of both **19** (> 50 kg) and **CMX990** (> 4 kg per batch) under conditions requiring no chromatographic separations (not shown in this disclosure). The drug substance **CMX990** has excellent handleability and is stable for >1 year at room temperature storage. It is crystalline in nature, and the structure was unambiguously assigned by x-ray crystallography (**Figure 3**) using starting material stereochemistries that are unlikely to invert during synthesis. PXRD (not shown) remains consistent during mechanical handling (such as micronization), and the material was readily amenable for the development of both simple aqueous suspension (e.g., in 0.5% methyl cellulose) or solid dosage forms. The stereochemistry of all four chiral centers of **CMX990** (C1, C6, C11, and C18; **Figure 3**) was found to be important for best activity, as confirmed with the synthesis (not shown) of all 16 possible stereoisomers of **CMX990**. P1-epimerization (**Figure 3** – at C6), often expected as the most likely to happen in a synthesis such as **Scheme 1**, led to >2x loss of potency (HeLa ACE-2 CoV-2 EC_90_ = 234 nM, HLM-Clint = 11.3 µL/min/mg) without impacting microsomal clearance. Inversion of P3-hydroxy group (**Figure 3** – at C18, HeLa ACE-2 CoV-2 EC_90_ = 196 nM, HLM-Clint = 23.7 µL/min/mg, F_PO(dog)_ = 55%) or the pyrrolidinone stereocenter (**Figure 3** – C1 epimer; HeLa ACE-2 CoV-2 EC_90_ = 150 nM, HLM-Clint = 13.9 µL/min/mg) was least critical, with the proline stereochemistry being the most critical (**Figure 3** – C11 epimer; HeLa ACE-2 CoV-2 EC_90_ = 6 µM, HLM-Clint = 31.0 µL/min/mg). Given the importance of ensuring chiral purity, significant effort was taken in developing manufacturing processes for **CMX990** to mitigate epimerization as well as to establish appropriate purification methods (particularly using alternative dissolution profiles / recrystallization). Target engagement of **CMX990** was validated in a biochemical assay for SARS-CoV-2 M^pro^, where it was demonstrated that the compound engages the enzyme as a reversible inhibitor (IC_50_ = 23.4 nM, **Table 2**). The reversible/irreversible character of the compound was assessed via different preincubation times of the inhibitor and the enzyme. In general, an irreversible inhibitor will increase its capacity to block the enzyme with increasingly longer incubation times in the presence of enzyme prior to addition of substrate. Conversely, a constant IC_50_ supports a reversible mechanism ^17^. Within the same assay, compound **3** (same warhead) was found to be a reversible inhibitor, while an analogous compound to **2** (aryloxymethyl ketone warhead) was found to be an irreversibly covalent inhibitor. **CMX990** was also screened in a pan-coronavirus reporter assay panel (data not shown), demonstrating the ability to potently inhibit up to 20 coronaviral protease types/variants including P132H (omicron), MERS, and HKU1(human), but not the negative control flavivirus system (ZIKV, no inhibition).

The *in vitro* activity of **CMX990** in a HeLa-ACE2 cell culture model of SARS-CoV-2 infection provided a mean antiviral EC_50/90_ of 37/90 nM, with no detectable cytotoxicity observed (CC_50_ > 40 µM). **CMX990** potently and reproducibly inhibited the replication of eight SARS-CoV-2 variants tested, including alpha, delta, and omicron (**Table 2**). It also showed activity versus other betacoronaviruses (e.g., HCoV-OC43) as well as alphacoronaviruses (e.g., HCoV-229E) in live virus assays, corroborating the findings in reporter assay data to designate **CMX990** as a pan-coronavirus inhibitor.

**Scheme 1:**
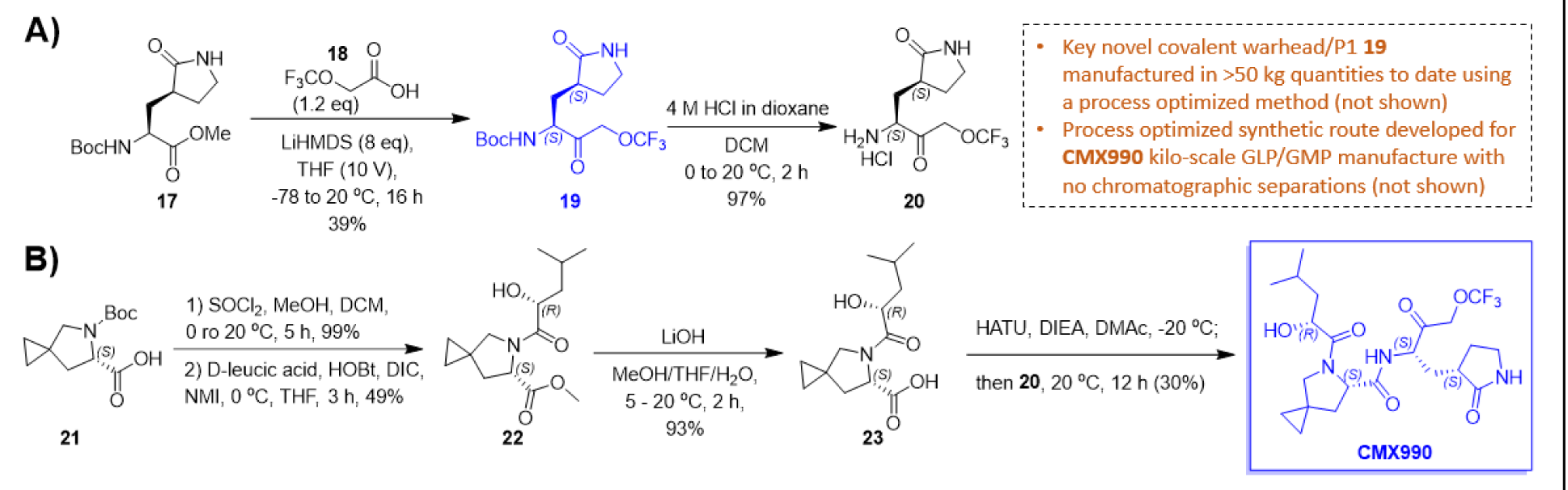
Synthesis of clinical candidate CMX990. Novel warhead-containing P1 fragment (**19**) can be manufactured at multi-kilo scales at >98% purity and stored long-term. The **CMX990** manufactured using this process is a white crystalline solid, readily amenable for both suspension formulation development as well as solid dosage form development. A) Synthesis of the novel trifluoromethoxymethyl ketone warhead bearing P1 unit (**19**). B) Core synthetic route for **CMX990**.

**Table 2:** Comparative performance of clinical candidate CMX990 vs. Nirmatrelvir.

|  | *Nirmatrelvir* | *CMX990* |
| --- | --- | --- |
| ***In Vitro Potency*** |  |  |
| SARS-CoV-2 M<sup>pro</sup> IC<sub>50</sub> [nM] | 28.1 | 23.4 |
| HeLa ACE2 CoV-2 EC<sub>50/90</sub> [nM] | 70 / 138 | 37 / 90 |
| ALI-HBEC CoV-2 EC<sub>50/90</sub> [nM] | 25.3 / 44.8 | 5.3 / 9.6 |
| HCT-8 hCoV-OC43 EC<sub>50/90</sub> [nM] | 112 / 418 | 135 / 712 |
| MCR5 hCoV-229E EC<sub>50/90</sub> [nM] | 353 / 798 | 25 / 146 |
| HeLa ACE2 CoV-2 *variants* EC<sub>90</sub> [nM]<br>alpha / delta / omicron | 242 / 196 / 117 | 63 / 35 / 37 |
| Uninfected Cytotoxicity CC<sub>50</sub> [μM]<br>HeLa-ACE2/HCT-8/MRC5/ALI HBEC | >40 / >30 / >30 / >3.3 | >40 / >30 / >30 / >3.3 |
| ***ADME Properties* [mu/rat/dog/cyno/human]** |  |  |
| Microsomal / hepatocyte stability<br>[CL<sub>int</sub>, μL/min/mg ; CL<sub>hep</sub>, μL/min/10<sup>6</sup>] | 32.8 / 2.3 (human) | 7.5 / 0.2 (human) |
| Human plasma stability [% at 2h] | >80% | >80% |
| PPB [%] | 73.6 / 58.6 / 98.8 / 70.3 / 54.8 | 48.1 / 88.8 / 63.6 / 62.3 / 56.3 |
| Microsomal protein binding [%] | 24.0 / 17.6 / 9.8 / 21.7 / 16.5 | 22.6 / 23.2 / 24.5 / 18.0 / 19.1 |
| B/P Ratio | 0.67 / 0.85 / 0.67 / 0.83 / 1.12 | 0.89 / 0.94 / 0.74 / 0.88 / 1.15 |
| P<sub>app(a->b/b->a)</sub> Caco-2 [10<sup>-6</sup> cm/s] | 0.9 / 9.0 | 0.6 / 9.3 |
| Aqueous Solubility | n.d. | 1.4-1.5 mg/mL |
| ***In Vivo Properties*<sup>a</sup> [mu/rat/dog/cyno]** |  |  |
| Plasma clearance<br>[IV PK CL<sub>p</sub> ; mL/min/kg] | 35.8 / 33.5 / 1.9 / 30.0 | 125 / 26.3 / 16.5 / 32.9 |
| IV PK V<sub>ss</sub> [L/kg] | 2.0 (Vd) / 1.0 / 0.57 / 1.5 | 1.2 / 2.7 / 1.1 / 1.2 |
| F<sub>po</sub> (%) | 17 / 44 / 69 / 3 | 14 / 12 / 53 / 1 |
| ***Target Selectivity and Tox Profiling*** |  |  |
| CEREP Screen (10 μM, 44 targets) | No hits | No hits |
| hERG channel (CHO cells) IC<sub>50</sub> | >10 μM | >10 μM |
| CYP inhibition IC<sub>50</sub><br>(1A2, 2C19, 2C9, 2D6, 3A4) | 3A4 = 46.4 μM,<br>rest > 50 μM | >50 μM (all) |
| Ames and MNT assay | Negative | Negative |
| Human peptidase selectivity panel | No significant engagement except below cathepsins |  |
| Cathepsin B / K / L / L2 / S IC<sub>50</sub> (μM) | >30 / 1.2 / >30 / 14.6 / 1.8 | 4.3 / 0.10 / 1.6 / 0.2 / 0.7 |

<sup>a</sup>further detailed in vivo data for **CMX990** in Table 5.

**Table 3:**
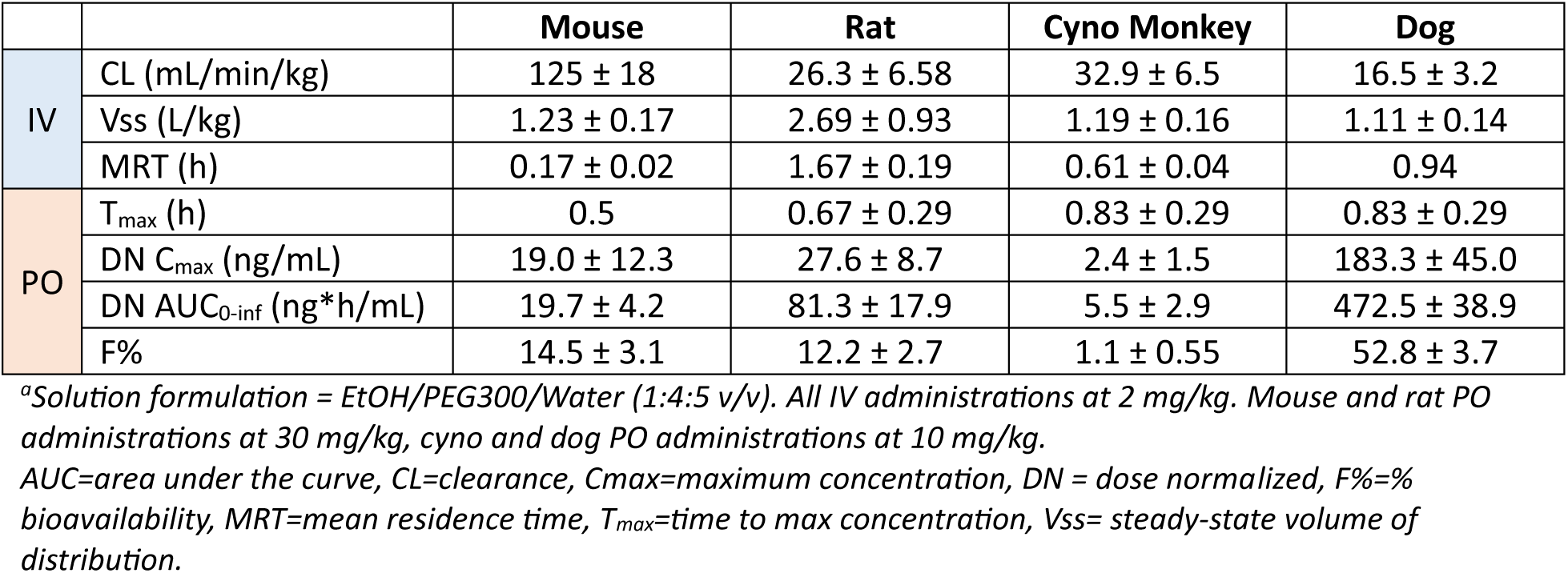
Pharmacokinetic parameters of CMX990 following a single IV or PO administration in solution formulation^a^.

|  |  | Mouse | Rat | Cyno Monkey | Dog |
| --- | --- | --- | --- | --- | --- |
| IV | CL (mL/min/kg) | 125 ± 18 | 26.3 ± 6.58 | 32.9 ± 6.5 | 16.5 ± 3.2 |
|  | V<sub>ss</sub> (L/kg) | 1.23 ± 0.17 | 2.69 ± 0.93 | 1.19 ± 0.16 | 1.11 ± 0.14 |
|  | MRT (h) | 0.17 ± 0.02 | 1.67 ± 0.19 | 0.61 ± 0.04 | 0.94 |
| PO | T<sub>max</sub> (h) | 0.5 | 0.67 ± 0.29 | 0.83 ± 0.29 | 0.83 ± 0.29 |
|  | DN C<sub>max</sub> (ng/mL) | 19.0 ± 12.3 | 27.6 ± 8.7 | 2.4 ± 1.5 | 183.3 ± 45.0 |
|  | DN AUC<sub>0-inf</sub> (ng*h/mL) | 19.7 ± 4.2 | 81.3 ± 17.9 | 5.5 ± 2.9 | 472.5 ± 38.9 |
|  | F% | 14.5 ± 3.1 | 12.2 ± 2.7 | 1.1 ± 0.55 | 52.8 ± 3.7 |

<sup>a</sup>Solution formulation = EtOH/PEG300/Water (1:4:5 v/v). All IV administrations at 2 mg/kg. Mouse and rat PO administrations at 30 mg/kg, cyno and dog PO administrations at 10 mg/kg.
AUC=area under the curve, CL=clearance, C<sub>max</sub>=maximum concentration, DN = dose normalized, F%= % bioavailability, MRT=mean residence time, T<sub>max</sub>=time to max concentration, V<sub>ss</sub>= steady-state volume of distribution.

As a more physiologically relevant model, SARS-CoV-2 replication was measured in normal primary human bronchial epithelial cells (HBECs). Air-liquid interface (ALI) cultures of HBECs recapitulate key properties of the airway epithelium, including cell polarization, mucus secretion, and cilia beating. **CMX990** performed 5x superior to nirmatrelvir in ALI-HBECs with a EC_50/90_ = 5.3/9.6 nM, having up to a 3-log reduction in viral load (72 h post-infection) when treated at 370 nM. No cytotoxicity was detected in any of these cell-based models.

**CMX990** has low to moderate plasma protein binding across species over the range of 0.5-20 µM as determined by equilibrium dialysis (**Table 2**). No meaningful concentration differences (i.e., >2-fold) in protein binding were observed in mouse, dog, monkey, and human plasma, with a concentration-dependent decrease in protein binding noted in rat at 20 µM. No meaningful species differences in protein binding were observed between species at 20 µM. The protein binding in human was comparable to mouse and dog, the two species selected for toxicity studies. **CMX990** (1 µM) had mean blood-to-plasma ratios (B/P) of 0.89, 0.94, 0.74, 0.88, and 1.15 in mouse, rat, dog, monkey, and human, respectively, suggesting no preferential partitioning into red blood cells in these species.

Metabolic turnover of **CMX990** was low in rat, dog, and human, with more rapid metabolism in mouse and monkey (**Table 2**). In microsomal stability studies conducted in the presence or absence of selective CYP450 inhibitors and in stability studies with individual recombinant CYP isozymes (CYP1A2, 2B6, 2C8, 2C9, 2C19, 2D6, 3A4), CYP3A4 was identified as the only enzyme contributing to the metabolism of **CMX990**. Excretion of the drug is anticipated to happen via both bile and urine. The route of elimination of **CMX990** was evaluated in bile duct cannulated rats. Following a single 3 mg/kg intravenous bolus dose, plasma, urine, and bile samples were collected up to 24 hours post-dose. The plasma half-life was short, most of the drug was eliminated by 7 hours post-dose, and ∼6.7% and 7.5% of unchanged **CMX990** was excreted to bile and urine, respectively. These data suggested that multiple clearance mechanisms are involved in the elimination of **CMX990**.

No critical off-target safety concerns were found for **CMX990** in a CEREP panel screen (Eurofins, 44 pharmacological targets, no individual target inhibition ≥ 50% at 10 µM concentration) or in a cardiac panel screen (including Nav1.5, Cav1.2, hERG). Peptidase panel inhibition data (Eurofins, at 10 µM) revealed no significant engagement of critical human peptidases (including calpain-1, caspases, chymase, chymotrypsin, DPP4, elastase 2, plasma Kallikrein, and plasmin) except some cathepsin isoforms. Some extent of *in vitro* cathepsin inhibition is observed in many of the comparative SARS CoV-2 peptidomimetic inhibitors we have studied to date including nirmatrelvir. Genotoxicity and mutagenicity concerns were also excluded via negative micronucleus (MNT) and Ames assay data (**Table 2**). Direct and time-dependent inhibition potential of **CMX990** on CYP1A2, CYP2C9, CYP2C19, CYP2D6, and CYP3A4 was investigated in human liver microsomes, where **CMX990** did not inhibit these major CYP isoforms (IC_50_ > 50 µM). There was no apparent shift in CYP3A4 IC_50_ (>50 µM) with a 30 min pre-incubation indicating **CMX990** is not a time-dependent inhibitor of CYP3A4.

**CMX990**’s pharmacokinetic profile was characterized by moderate plasma clearance values in dog and rat, with higher clearance in monkey and mouse, following 2 mg/kg intravenous doses (**Table 3**). The steady-state volume of distribution was moderate in all species (Vss 1.0–2.3 L/kg). Plasma elimination half-life after IV dosing for **CMX990** was < 1 hr in both mouse and monkey, with values of 1.2–1.9 and 2.0 hours in rat and dog, respectively. The compound was rapidly absorbed, with T_max_ values ranging between 0.33 to 1.7 hr across species. Oral bioavailability was low in monkey (1%), moderate in mouse and rat (14-22%) and high in dog (53%; **Table 3**), which was similar to the preclinical profile seen for nirmatrelvir in comparative in-house studies (**Table 2**). Low oral bioavailability in monkey for such peptidomimetics has been presented in prior literature ^18^. *In vivo* hepatic clearance predicted from *in vitro* intrinsic clearance (using scaling factors and well-stirred liver model) showed good correlation with measured *in vivo* clearance trends in preclinical animals (**Tables 2 and 3**).

Based on this excellent *in vitro* to *in vivo* correlation (IVIVc) of clearance from other species, human clearance for **CMX990** was predicted to be very low (predicted CL_h_ < 5.9 [microsomal], < 3.0 [hepatocytes]). This led to human-equivalent dose projections (oral, < 500 mg/day) of once- or twice- a- day with no ritonavir boost required to cover efficacious C_trough_ exposure, significantly differentiating **CMX990** from nirmatrelvir. A ritonavir boosted study (not shown) done in dogs showed no significant advantage towards the pharmacokinetic profile of **CMX990** versus monotherapy.

Multiple mono-, di-, and tri-oxidative metabolites were identified (not shown) following incubations in microsomes (8 metabolites) and hepatocytes (15 metabolites) across species. No human unique metabolites were identified in either microsome or hepatocyte incubations, and all human metabolites were present in the species selected for toxicology studies (e.g., mouse, dog, or both). The intended clinical route of administration for **CMX990** was decided to be oral administration; therefore, all the toxicology studies were conducted by oral gavage.

Several additional studies (not shown) explored the effect of formulation, dose, and dosing frequency on **CMX990** exposures following oral administration in CD-1 mice and beagle dogs. In mice, dose-normalized AUC values were comparable between the 30 mg/kg solution formulation and the 35, 50, 100, 300 and 500 mg/kg BID doses of a suspension of micronized API (active pharmaceutic ingredient), with greater dose proportional values following the 50 mg/kg BID dose. In dogs, exposures from aqueous suspensions of micronized API (D_90_ < 50 µM) were higher than those obtained from aqueous suspensions of non-micronized drug, with BID dosing providing greater than proportional increases in AUC when compared to the same single daily dose. Taken together, BID dosing of the micronized API as a suspension in 0.5% methylcellulose (MC) was selected as the formulation for repeat-dose toxicology studies.

The general toxicity profile of **CMX990** was assessed in CD-1 mice and beagle dogs. **CMX990** was administered by oral gavage to CD-1 mice and beagle dogs at dosages of up to 1000 mg/kg/day (500 mg/kg/dose, BID, 8 hours apart) for five days with a 14-day recovery period. **CMX990** was well-tolerated in both species. In mice, non-adverse microscopic findings only included sporadic **CMX990**-related minimal to mild increased single-cell necrosis of hepatocytes without associated changes in serum liver enzymes. In dogs, liver was the primary target organ with microscopic findings of adverse centrilobular hepatocellular degeneration/necrosis that was accompanied by increases in serum liver enzymes. Additional non-adverse findings in dog also included epithelial single-cell necrosis and mononuclear cell infiltration in gall bladder, decreased cellularity in thymus, decreases in reticulocytes and increases in clotting times. These findings were generally monitorable and were either completely reversible or were in the process of resolution (liver) at the end of the recovery phase. The no observed adverse effect levels (NOAELs) in mice and dogs are considered to be 1000 mg/kg/day (500 mg/kg/dose, BID) and 150 mg/kg/day (75 mg/kg/dose, BID), respectively for 5-day administration.

## CONCLUSION

Inhibition of the highly conserved viral main protease (M^pro^) is one of the best accepted means to develop antivirals to coronaviruses. SARS-CoV-2 M^pro^ has been well validated as a target with an FDA-approved drug now on the market, as well as from a large body of research that has evolved over the past three years. Currently available therapies for SARS-CoV-2 infection have an array of disadvantages including DDI and other toxicology concerns, global accessibility, as well as suboptimal potencies of target engagement. Particularly in the view of future pandemic preparedness, it is imperative that further improvements are made to develop better and/or alternative pharmaceuticals that can tackle this drug target, which has no human homolog. The key breakthrough of the described program to address this need was based on a unique unprecedented cysteine protease-engaging warhead that enabled low nanomolar antiviral potencies.

**CMX990** features a novel covalent warhead, excellent antiviral properties (SARS CoV-2 EC_90_ [ALI HBEC] = 10 nM), good oral bioavailability and tolerability in preclinical species, excellent human *in vitro* metabolic clearance (Pred. CL_h_ < 6 mL/min/kg), and HED predictions (once or twice a day dosing) supporting unboosted oral therapy. With this promising background and the support of a large multidisciplinary team, **CMX990** progressed to a Phase 1 clinical trial in ∼10 months from its first synthesis.

## EXPERIMENTAL SECTION

### Materials and Methods

All reagents and solvents were purchased from commercial sources and used without further purification. Flash column chromatography was performed using silica gel (200−300 mesh). All reactions were monitored by TLC (pre-coated EMD silica gel 60 F254 TLC aluminum sheets and visualized with a UV lamp or appropriate stains) and/or LCMS (Waters Acquity UPLC system, 2 or 4 min run of a 10-90% mobile phase gradient of acetonitrile in water [+0.1% formic acid]). NMR spectra were obtained on Bruker AV400 or AV500 instruments, and data was analyzed using the MestReNova NMR software (Mestrelab Research S. L.). Chemical shifts (δ) are expressed in ppm and are internally referenced for ^1^H NMR (CHCl_3_ 7.26 ppm, DMSO-*d_6_* 2.50 ppm) and ^13^C NMR (CDCl_3_ 77.16 ppm, DMSO-*d_6_* 39.52 ppm). X-ray data was collected at room temperature on a Bruker D8 QUEST instrument with an IμS Mo microfocus source (λ = 0.7107 Å) and a PHOTON-III detector.

NMR spectra for the final proline containing derivatives commonly exist as rotamer populations in DMSO-*d*_6_. The activated ketone in the novel warhead creates a transient reversible-equilibrium under acid modified (f.ex., TFA, formic acid) aqueous chromatography conditions, as represented by the appearance of a leading non-baseline separated second (minor) peak in the chromatogram. Occasionally, some compounds (f.ex. Compound **10** with truncated P2-cap) also show similar interactions/equilibrium in NMR, including interactions of the P2-proline nitrogen with the warhead ketone. The isolation of individual components of these non-baseline separated material, and re-injection results in the same equilibrium appearance. Hence, when doing chromatographic separations under reversed-phase conditions containing acidic modifiers, the entirety of this peak (leading minor-peak + the later-eluting major peak) should be isolated as a single peak. Alternatively, reversed-phase separation can be carried out without the presence of an acidic modifier, with a slight compromise on sharpness of the peak.

#### Chemistry

All compounds with the novel trifluoromethoxymethyl warhead were synthesized according to the representative procedure outlined for **CMX990** below. All compounds are >95% pure by HPLC and/or NMR analyses.

#### *tert*-butyl ((*S*)-3-oxo-1-((*S*)-2-oxopyrrolidin-3-yl)-4-(trifluoromethoxy)butan-2-yl)carbamate (19)

To a stirred solution of 2-(trifluoromethoxy)acetic acid **18** (8.45 g, 58.67 mmol, 1.2 eq) in THF (90 mL), lithium bis(trimethylsilyl)amide solution (1 M in THF, 391.16 mL, 391.16 mmol, 8.1 eq) was added and the resulting mixture was stirred at −78°C for 1 h. Methyl (*S*)-2-((*tert*-butoxycarbonyl)amino)-3-((*S*)-2-oxopyrrolidin-3-yl)propanoate (**17**) (14.0 g, 48.89 mmol, 1.0 eq) in THF (50 mL) was slowly added to the reaction mixture at −78 °C and the reaction mixture was stirred at −78 °C for 3 h and at room temperature for 16 h. After completion of the reaction, the reaction mixture was quenched with AcOH (28 mL) in THF (126 mL) at 0 °C and diluted with water (500 mL). The resulting mixture was filtered through celite and washed with EtOAc (500 mL), and filtrate was extracted with EtOAc (3 × 500 mL). The combined organic layer was dried over anhydrous Na_2_SO_4_, filtered, evaporated under reduced pressure to get crude compound. The resulting crude compound was purified by davisil grade silica gel column chromatography using 20% ethyl acetate in hexane as the eluent to afford *tert*-butyl ((*S*)-3-oxo-1-((*S*)-2-oxopyrrolidin-3-yl)-4-(trifluoromethoxy)butan-2-yl)carbamate (**19**, 6.79 g, 19.2 mmol, 39%) as a pale brown solid. [*TLC system:* EtOAc: pet-ether (7:3); *R_f_* value: 0.4]. LCMS *m/z* 355.37 (M+1); ^1^H NMR (400 MHz, DMSO-*d_6_*) δ 7.66 (s, 1H), 7.52 (dd, *J* = 7.2 Hz, 18 Hz, 1H), 4.98 (s, 2H), 4.15-4.09 (m, 1H), 3.19-3.12 (m, 2H), 2.50-2.13 (m, 2H), 1.91-1.84 (m, 1H), 1.68-1.57 (m, 2H), 1.39 (s, 9H).

#### (3*S*)-3-[(2*S*)-2-amino-3-oxo-4-(trifluoromethoxy)butyl]pyrrolidin-2-one hydrochloride (20)

To a stirred solution of *tert*-butyl *N*-[(2*S*)-3-oxo-1-[(3*S*)-2-oxopyrrolidin-3-yl]-4-(trifluoromethoxy)butan-2-yl]carbamate (**19**; 20 g, 56.5 mmol, 1.0 eq) in DCM (300 mL) was added HCl (4 M in dioxane, 34.30 mL, 20 eq) dropwise at 0 °C under nitrogen atmosphere. The resulting mixture was stirred for additional 1h at room temperature. The resulting mixture was concentrated under reduced pressure. The residue was purified by trituration with Et_2_O (150mL) to afford (3*S*)-3-[(2*S*)-2-amino-3-oxo-4-(trifluoromethoxy)butyl]pyrrolidin-2-one hydrochloride as an off-white solid (**20**; 16 g, 55.0 mmol, 97% yield). LCMS *m/z* = 255 (M+1).

#### Methyl (*S*)-5-azaspiro [2.4] heptane-6-carboxylate hydrochloride (22)

To a stirred solution of the starting proline **21** (150 g, 622 mmol, 1.0 eq) in MeOH (2 L) was slowly added SOCl_2_ (221.86 g, 1.86 mol, 3 eq) and DMF (1.36 g, 18.65 mmol, 0.03 eq) dropwise at 0°C under nitrogen atmosphere. The resulting mixture was stirred for 5 h at room temperature. After the reaction was completed, the mixture was concentrated under reduced pressure to afford methyl (*S*)-5-azaspiro [2.4] heptane-6-carboxylate hydrochloride **22** (120 g, 615 mmol, 99%) as a light-yellow oil.

#### Methyl (*S*)-5-((*R*)-2-hydroxy-4-methylpentanoyl)-5-azaspiro [2.4] heptane-6-carboxylate (23)

To a stirred mixture of D-Leucic acid (100 g, 522 mmol, 1.0 eq) and (*R*)-2-hydroxy-4-methylpentanoic acid **22** (75.85 g, 574 mmol, 1.1 eq) in THF (3 L) were slowly added DIC (79.02 g, 626 mmol, 1.2 eq) and 1-methyl-1*H*-imidazole (171.36 g, 2.09 mol, 4 eq) in portions at 0 °C. The resulting mixture was stirred for 10 min at 0°C. Then a solution of HOBt (84.61 g, 626 mmol, 1.2 eq) in THF (200 mL) was slowly added dropwise over 30 min. The resulting mixture was stirred for 3 h at 0°C, filtered, and the filtrate concentrated under reduced pressure. This residue was purified by silica gel column chromatography, eluted with PE: THF (5:1), to afford partially pure **23** (69 g, 256 mmol, 49%) as a white solid (with intramolecularly trans-esterified compound as an impurity).

#### (*S*)-5-((*R*)-2-hydroxy-4-methylpentanoyl)-5-azaspiro [2.4]heptane-6-carboxylic acid (24)

A solution of **23** (69 g, 256 mmol, 1.0 eq) in THF (700 mL) was stirred at 0 °C. LiOH (7.36 g, 307 mmol, 1.2 eq) in H_2_O (200 mL) was added to this dropwise over 20 min while maintaining the reaction mixture temperature at 0°C. The resulting mixture was stirred for 3 h at room temperature, acidified to pH = 3 with 50 mL HCl (6 N) at 0 °C, and then extracted with EtOAc (3 × 500 mL). The combined organic layers were washed with brine (250 mL), dried over with 250 g anhydrous Na_2_SO_4_, filtered, and concentrated *in vacuo* to afford **24** (61 g, 239 mmol, 93%) as a white solid.

#### (*S*)-5-((*R*)-2-hydroxy-4-methylpentanoyl)-*N*-((*S*)-3-oxo-1-((*S*)-2-oxopyrrolidin-3-yl)-4-(trifluoromethoxy)butan-2-yl)-5-azaspiro[2.4]heptane-6-carboxamide (CMX990)

To a stirred solution of acid **24** (61 g, 239 mmol, 1.0 eq) and amine hydrochloride **20** (85.97 g, 263 mmol, 1.1 eq) in DMAc (1.2 L) was added HATU (109.02 g, 287 mmol, 1.2 eq). The mixture was stirred for 10 min at −20°C in an ethanol-ice bath, and DIEA (123.52 g, 956 mmol, 4.0 eq) diluted with 250 ml DMAc was slowly added. The resulting mixture was stirred for 12 h at room temperature, then diluted with water (1.5 L), and extracted with EtOAc (2 × 2.0 L). The combined organic layers were washed with brine (500 mL), dried over with anhydrous Na_2_SO_4_, filtered, and concentrated *in vacuo*. The residue was taken up in EtOAc (500 mL) and washed with 1 M hydrochloric acid (3 x 250 mL). The water layers were extracted with EtOAc (2 × 500 mL), combined organic layers dried over with anhydrous Na_2_SO_4,_ filtered, and concentrated *in vacuo*. The resultant residue was purified by silica gel column chromatography, eluted with PE: THF=1:1 to afford 27 g (SFC: 94% purity, batch A) and 19 g (SFC: 91% purity, batch B)

**CMX990**. 19 g (SFC: 91% purity, batch B) was purified by SFC to give 12 g material (SFC: 98% purity, batch C). Batch A and C were combined and slurried with Et_2_O (400 mL) to give **CMX990** (35.8 g, 72.8 mmol, 30%) as a white crystalline solid.

SFC separation condition: Column: CHIRAL ART Cellulose-SC, 3*25 cm, 5 μm; Mobile Phase A: CO2, Mobile Phase B: IPA: MeCN = 1: 1; Flow rate: 80 mL/min; Gradient: isocratic 25% B; Column Temperature (℃): 35; Back Pressure(bar): 100; detection wavelength: 220 nm; Sample Solvent: IPA: MeCN = 2:1; Injection Volume: 5 mL (x 110 runs)

HRMS *m/z* calcd. for C_22_H_33_F_3_N_3_O_6_ ^+^ [M+H]^+^ 492.2316, found 492.4428; ^1^H NMR (400 MHz, DMSO-*d*_6_) δ 8.67 (br d, *J* = 7.0 Hz, 0.2H), 8.43 (br d, *J* = 7.7 Hz, 0.7H), 7.73 (br s, 0.2H), 7.67 (br s, 0.7H), 5.10 (d, *J* = 7.3 Hz, 0.2H), 5.02 (d, *J* = 17.4 Hz, 1.2H, some overlap), 4.88 (d, *J* = 17.3 Hz, 1H), 4.60 (d, *J* = 7.3 Hz, 0.7H), 4.35 (m, 1.8H), 4.11 (m, 0.8H), 3.91 (m, 0.2H), 3.58 – 3.46 (m, 1.8H), 3.22 – 3.06 (m, 2.2H), 2.30 – 2.06 (m, 3H), 2.03 – 1.89 (m, 1H), 1.84 – 1.54 (m, 4H), 1.51 – 1.18 (m, 2H), 0.88 (d, *J* = 6.8 Hz, 2.3H), 0.87 (d, *J* = 6.5 Hz, 2.3H), 0.88 (d, *J* = 6.7 Hz, 0.7H), 0.87 (d, *J* = 6.5 Hz, 0.7H), 0.66 – 0.37 (m, 4H); ^19^F NMR (376 MHz, CDCl_3_) δ −60.86; ^13^C NMR (101 MHz, CDCl_3_) δ 200.80, 180.44, 174.63, 172.76, 121.63 (q, *J* = 256.3 Hz), 69.04 (q, *J* = 2.9 Hz), 61.50, 56.41, 54.10, 43.17, 40.94, 38.76, 37.26, 31.28, 28.89, 24.62, 23.72, 21.65, 21.09, 13.69, 7.21, 1.13; Melting point = 170.2 – 172.6 °C; Anal. calcd. for C_22_H_32_F_3_N_3_O_6_: C, 53.76; H, 6.56; N, 8.55 Found: C, 53.34; H, 6.60; N, 8.41.

## AUTHOR INFORMATION

### Notes

Authors declare no competing financial interests.

## ACKNOWLEDGEMENTS

This work was supported by grants from the Bill and Melinda Gates Foundation (INV-028691, COVID-19 CTA GPP). Authors would like to thank Sreehari Babu Putchakayala (Aragen Life Sciences) for assistance in formulations development work, Dr. Jason Chen and his team at the Scripps Research Automated Synthesis facility for assistance with chiral purifications/characterization, Calibr compound management team and administrative teams, Dr. Kit Bonin (Calibr) and Dr. Geneva Hargis (Calibr) for manuscript preparation assistance, and the diverse team of subject-matter-experts engaged by the Bill and Melinda Gates Foundation to guide and accelerate this program in to Ph1 clinical studies.

## ABBREVIATIONS

3CL^pro^: 3C-like protease
ACE-2: angiotensin-converting enzyme-2
ALI: air-liquid interfaces
CL: clearance
CLhep: clearance in hepatocytes
Clint: intrinsic clearance
Clp: plasma clearance
CL^pro^: 3C-like protease
C_max_: maximum concentration
C_trough_: concentration before next dose
F: bioavailability
DN: dose normalized
HBEC: human bronchial epithelial cells
HED: human equivalent dose
HKU1: human coronavirus 1
HLM: human liver microsomes
hPPB: human plasma protein binding
IVIVc: in vitro to in vivo correlation
logD: logarithm of partition coefficient for ionizable compounds
MC: methylcellulose
MERS: Middle Eastern respiratory syndrome
met-ID: metabolite identification
MNT: micronucleus
M^pro^: main protease (i.e., 3CL^pro^)
MRT: mean residence time
Mu: mouse
n.c.: not calculated
n.d.: not determined
P1: amino acid 1, recognition element with a lactam side chain
P132H: omicron variant of SARS-CoV-2
P2: amino acid 2
P3: a N-Cap or amino acid 3
P_app_: apparent permeability coefficient;PL^pro^, papain-like protease
ReFRAME: Repurposing, Focused Rescue, and Accelerated Medchem library
SARS-CoV-2: severe acute respiratory syndrome coronavirus 2
T_max_: time to peak drug concentration
V_ss_: steady state volume of distribution
WH: warhead
ZIKV: Zika virus

---

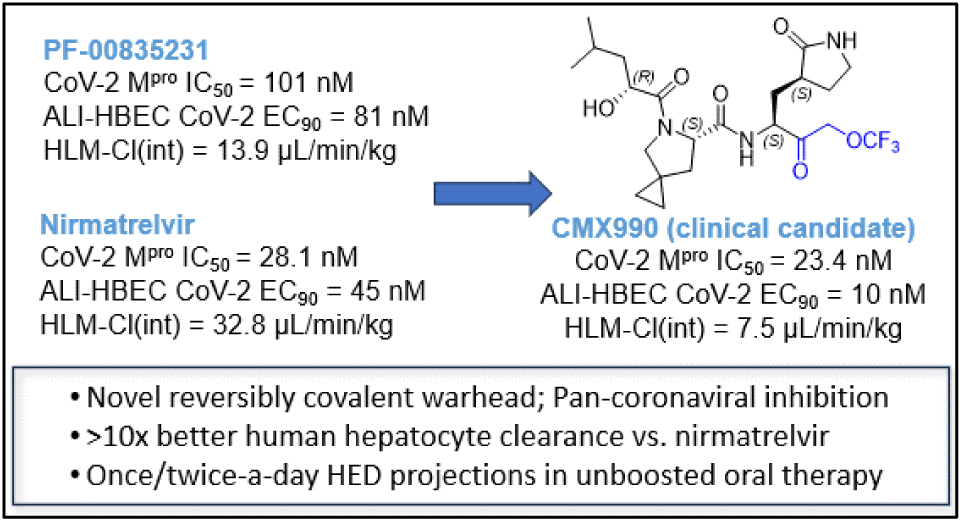
Table of Contents Graphic:

